# The molecular basis of ubiquitin-specific protease 8 autoinhibition by the WW-like domain

**DOI:** 10.1101/2021.05.31.446389

**Authors:** Keijun Kakihara, Kengo Asamizu, Kei Moritsugu, Masahide Kubo, Tetsuya Kitaguchi, Akinori Endo, Akinori Kidera, Mitsunori Ikeguchi, Akira Kato, Masayuki Komada, Toshiaki Fukushima

## Abstract

Ubiquitin-specific protease 8 (USP8) is a deubiquitinating enzyme involved in multiple membrane trafficking pathways. The enzyme activity is inhibited by binding to 14-3-3 proteins, and mutations of the 14-3-3 binding motif in USP8 are related to Cushing’s disease. However, the molecular basis of USP8 enzyme activity regulation remains unclear. Here, we identified amino acids 645–684 of USP8 as an autoinhibitory region, which our pull-down and single-molecule FRET assay results suggested interacts with the catalytic USP domain. *In silico* modelling indicated that the region forms a WW-like domain structure, plugs the catalytic cleft, and narrows the entrance to the ubiquitin-binding pocket. Furthermore, 14-3-3 was found to inhibit USP8 enzyme activity partly by enhancing the interaction between the WW-like and USP domains. These findings provide the molecular basis of USP8 autoinhibition via the WW-like domain. Moreover, they suggest that the release of autoinhibition may underlie Cushing’s disease caused by USP8 mutations.

---

Ubiquitin-specific protease 8 (USP8), a member of the USP family of deubiquitinases, regulates multiple membrane trafficking pathways and membrane fusion/fission events. It deubiquitinates endosomal membrane proteins, including epidermal growth factor receptor (EGFR), which enhances their recycling to the plasma membrane and/or inhibits their lysosomal degradation [1–8]. USP8 also maintains the protein levels and/or functions of endocytosis machinery components including Eps15, HRS, STAM1/2 and CHMP proteins [9–12]. In addition, USP8 regulates autophagy and mitophagy by deubiquitinating EPG5, p62 and Parkin [13–15]; it suppresses the endoplasmic reticulum export of procollagen by deubiquitinating COPII protein Sec31 [16]; and it enables cytokinesis, probably by deubiquitinating VAMP8, a SNARE required for vesicle fusion in cytokinesis [17]. Given the many functions of USP8, it is important that the regulatory mechanism of its enzyme activity be elucidated; however, limited mechanistic data has been reported to date.

Similar to other USPs, USP8 harbours a catalytic USP domain consisting of three subdomains termed the fingers, palm and thumb [18, 19] (Fig. 1**a**). These subdomains cooperatively form the ubiquitin-binding pocket, which recognises the globular core of the distal ubiquitin molecule. A deep cleft between the palm and thumb subdomains functions as a catalytic cleft at which the catalytic triad (Cys, His and Asp/Asn residues) attacks the isopeptide bond between the distal ubiquitin C-terminal tail and proximal ubiquitin (or substrate protein). The crystal structure of the USP8 catalytic domain in its apo-form shows its unique features [20]; two loops, namely blocking loops 1 and 2 (BL1 and BL2), which are utilised for ubiquitin recognition in other USPs, are positioned in a closed conformation. Furthermore, the fingers subdomain is tightened inwardly, making the ubiquitin-binding pocket too narrow to capture ubiquitin. Despite these unfavourable features, the USP domain effectively catalyses the deubiquitination reaction [20], implying that substrate-induced conformational change occurs.

**Fig. 1.**
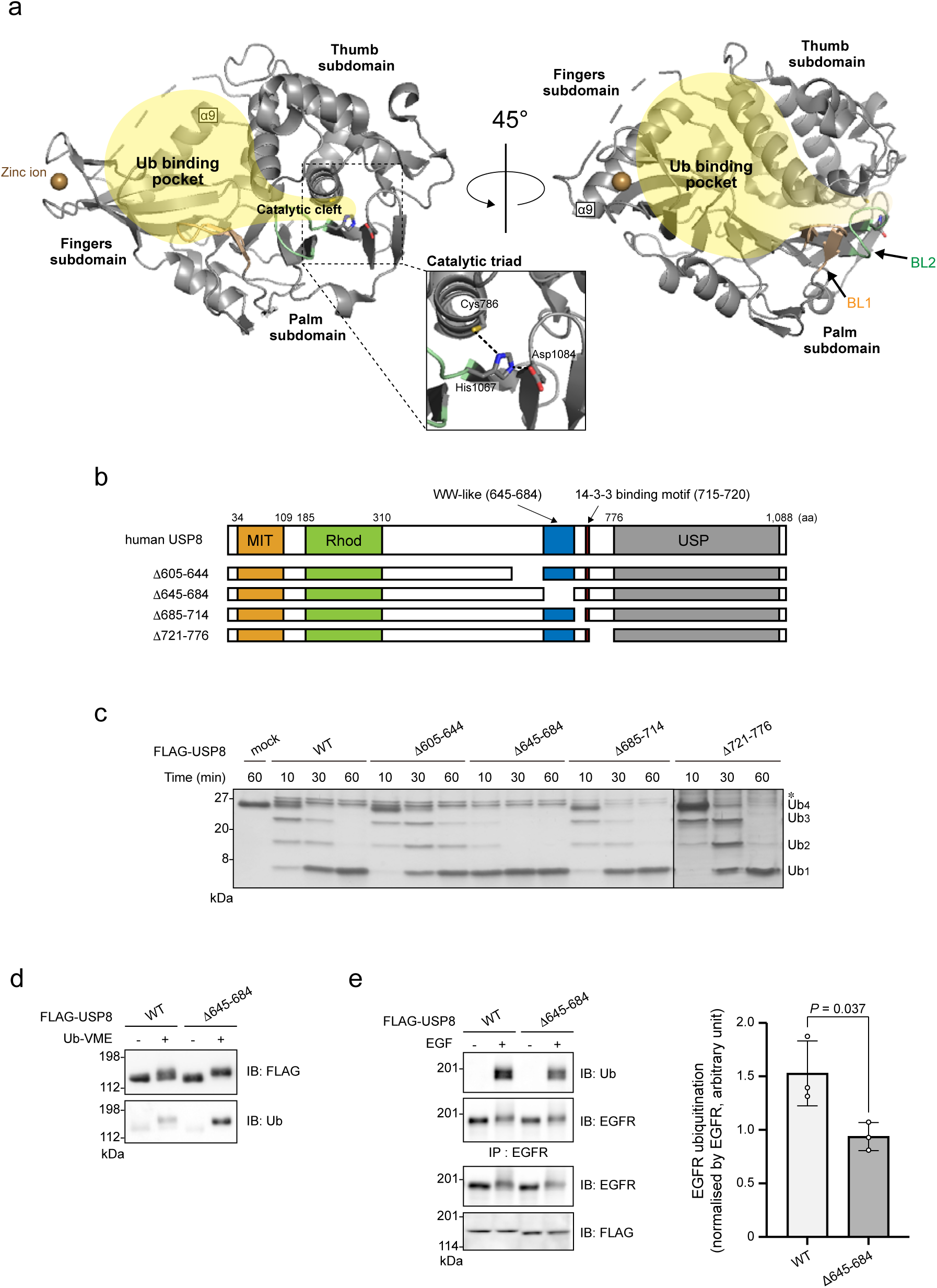
Effects of deleting USP8 amino acids 645–684 on its activity. **(a)** Crystal structure of the USP domain (aa 762–1109; PDB ID: 2GFO). Two representations are rotated by 45° around the Y-axis. The fingers, palm and thumb subdomains are indicated. The catalytic triad formed by Cys^786^, His^1067^ and Asp^1084^ is shown in the enlarged view. The ubiquitin-binding pocket and catalytic cleft are indicated in yellow. The upper-left broken loop between Asp^888^ and Asn^898^ is non-structural. BL1, blocking loop 1; BL2, blocking loop 2. **(b)** Schematic structures of human USP8 and the truncated mutants used in c: MIT, MIT domain; Rhod, rhodanese like domain; WW-like, WW-like domain identified in this study; USP, USP domain. **(c)** Deubiquitination activities of the USP8 mutants. HEK293T cells expressing FLAG-tagged USP8 mutants were lysed. Anti-FLAG immunoprecipitates were subjected to an *in vitro* deubiquitination assay using a Lys63-linked ubiquitin tetramer, and then to SDS-PAGE and silver staining. *, co-purified protein(s) with USP8. **(d)** Ubiquitin-vinyl methyl ester (Ub-VME)-labelling of USP8^Δ645–684^. Immunoprecipitated USP8 (WT or Δ645–684) were labelled and subjected to immunoblotting (IB). **(e)** EGFR ubiquitination levels in cells expressing USP8^Δ645–684^. HeLa cells stably expressing USP8 (WT or Δ645–684) were treated with or without EGF. The lysates were subjected to immunoprecipitation (IP) and immunoblotting. Right graph shows EGFR ubiquitination levels normalised by EGFR in the immunoprecipitates (means ± standard deviations of three independent experiments). Statistical significance was determined by Student’s *t*-test. See Supplementary Fig. 1.

Some USPs are activated by specific interacting proteins that bind to catalytic USP domains [21–25] or other domains [26–28]. USP8, which has a phosphorylation-dependent 14-3-3 binding motif (Fig. 1**b**), is inhibited by binding to 14-3-3 [29]. In the mitotic M-phase, when USP8 positively regulates cytokinesis, it dissociates from 14-3-3 and shows increased enzyme activity [29]. On the other hand, in corticotroph adenomas of Cushing’s disease, ∼50% of tumour samples show somatic mutations of the *USP8* gene that cause deletion or substitution of amino acids around the 14-3-3 binding motif [30, 31]. Cushing’s disease begins with hypersecretion of adrenocorticotropic hormone (ACTH) from corticotrophs, followed by excess adrenal glucocorticoid. In cultured corticotrophs, exogenous expression of USP8 lacking functional 14-3-3 binding motifs effectively enhances ACTH secretion [30], suggesting that 14-3-3 binding motif dysfunction causes this disease. Thus, 14-3-3 plays important roles in regulating USP8 under physiological and pathological situations. However, the molecular basis of 14-3-3-dependent inhibition of USP8 enzyme activity remains unclear.

This study was undertaken to elucidate the regulatory mechanism of USP8 enzyme activity. Here, we identified a novel WW-like domain in USP8 as an autoinhibitory region. This WW-like domain and ubiquitin were found to competitively bind to the ubiquitin-binding pocket of USP8. In addition, 14-3-3 was shown to inhibit USP8 enzyme activity partly by enhancing the interaction between the WW-like domain and catalytic domain.

## Results

### USP8 has an autoinhibitory region

We generated a series of USP8 deletion mutants (Fig. 1**b**; Supplementary Fig. 1**a**) and evaluated their deubiquitinating activities *in vitro*. In Supplementary Fig. 1**b**, a mutant lacking amino acids (aa) 414–714 (USP8^Δ414–714^) and USP8^Δ605–714^ deubiquitinated ubiquitin chains faster than wild-type USP8 (USP8^WT^) and other mutants, suggesting that aa 605–714 contains autoinhibitory region(s) that decreases USP8 deubiquitinating activity. Further mutational analyses determined aa 645–684 as a putative autoinhibitory region (Fig. 1**c**).

Next, we tested the labelling efficiency of USP8^WT^ or USP8^Δ645–684^ with ubiquitin-vinyl methyl ester (Ub-VME), which can be irreversibly conjugated to Cys residue of the catalytic triad [32]. Ub-VME was able to label USP8^Δ645–684^ more effectively than it labelled USP8^WT^ (Fig. 1**d**), implying that aa 645–684 hinders ubiquitin recognition by the USP domain.

We then examined the effects of deleting aa 645–684 on the ubiquitination levels of USP8 substrates in cells. EGF-induced ubiquitination of EGFR, a representative USP8 substrate, was lower in HeLa cells stably expressing USP8^Δ645–684^ than in cells expressing USP8^WT^ (Fig. 1**e**; Supplementary Fig. 1**c**). We also examined ubiquitination of other USP8 substrates, i.e. Parkin [15], STAM1 [10] and STAM2 [2], using HEK293T cells overexpressing USP8^WT^ or USP8^Δ645–684^. The ubiquitination levels of these substrates were decreased by USP8^WT^ overexpression but were decreased more strongly by USP8^Δ645–684^ overexpression (Supplementary Figs. 1**d–f**). These results suggest that aa 645–684 decreases USP8 deubiquitinating activity in cells. This region seems not to regulate intracellular localisation of USP8, however, because USP8^Δ645–684^ showed a similar staining pattern to that of USP8^WT^ (Supplementary Fig. 1**g**). Taking together with *in vitro* data (Figs. 1**b-d**), we concluded that aa 645–684 functions as an autoinhibitory region.

### The autoinhibitory region is classified as an atypical WW domain

The amino acid sequence of the autoinhibitory region is well conserved among vertebrates (Fig. 2**a**) and shows sequence similarity to the MAGI1 WW domain (aa 359–392) (Fig. 2**b**). Typical WW domains have two conserved tryptophan residues; however, some WW domains in MAGI1, SAV1 and WWOX have only the first residue (Fig. 2**b**), although these atypical WW domains show conformations that are similar to those of typical WW domains [33]. The autoinhibitory region of USP8 has only the first tryptophan residue (Trp^655^), i.e. it shares a common feature with atypical WW domains.

**Fig. 2.**
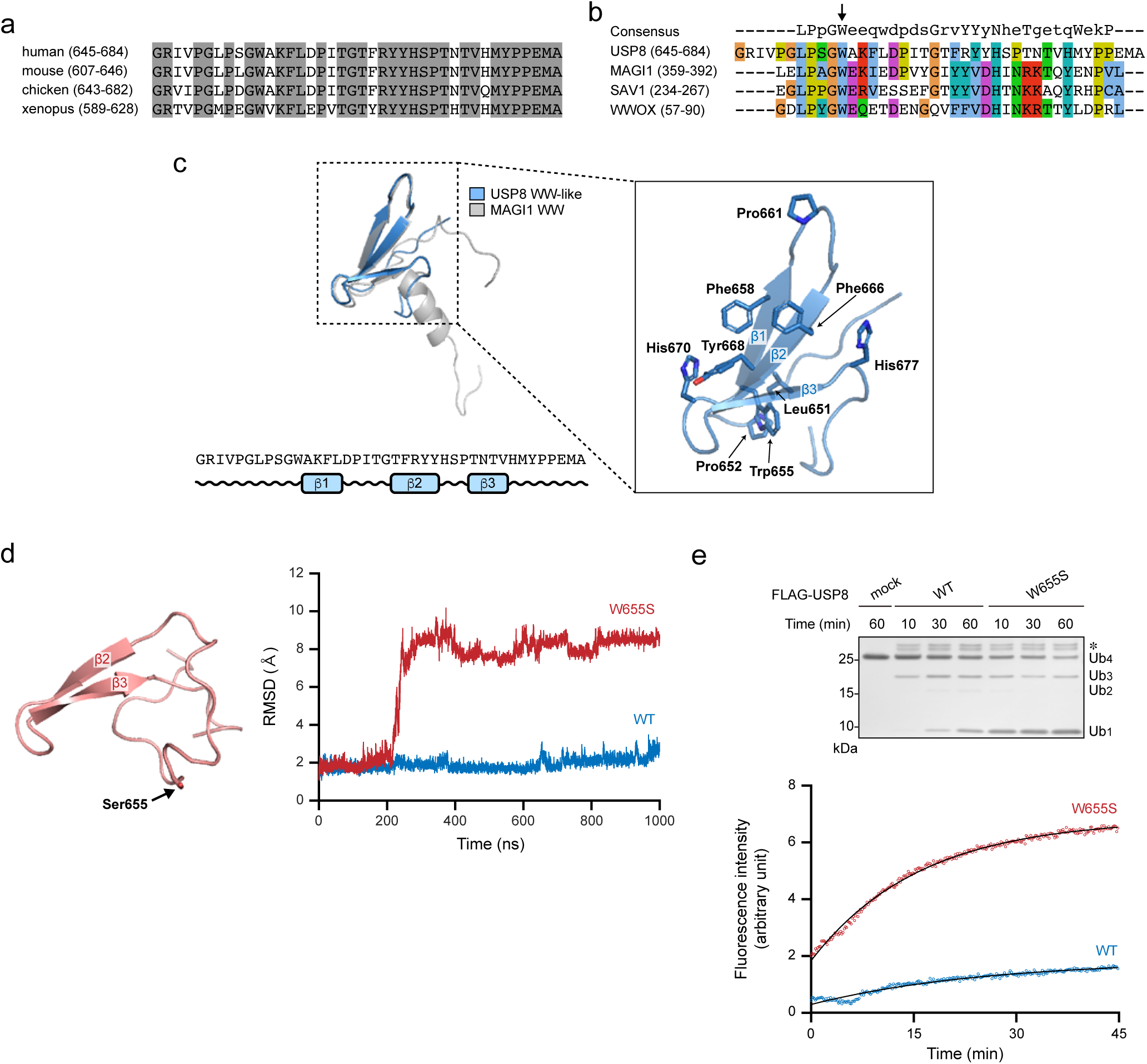
Structural modelling of the USP8 autoinhibitory region. **(a)** The amino acid sequence conservation among vertebrates. Grey, conserved residues. **(b)** The sequence similarity to the atypical WW domains of MAGI1, SAV1 and WWOX. The arrow indicates the conserved N-terminal tryptophan. The consensus sequence of WW domain is also shown. **(c)** A structural model of the WW-like domain. Left, superposition of the predicted model (blue) and MAGI1 WW domain (grey), as well as the secondary structure based on the model. Right, magnification of the hydrophobic surface formed by three anti-parallel β-sheets. **(d)** Molecular dynamics simulations of the predicted model and Trp^655^ to Ser (W655S) mutant. Left, the simulation structure of the W655S mutant. Right, the root-mean-square deviations (RMSD) from the predicted structure as a function of simulation time. Blue, WT. Red, W655S mutant. **(e)** Deubiquitination activities of USP8^W655S^. Top, ubiquitin chain cleavage assay by the similar methods to those described in Fig. 1c. *, co-purified protein(s) with USP8. Bottom, ubiquitin-AMC assay. HEK293 cells expressing USP8^W655S^ were lysed. Anti-FLAG immunoprecipitates were subjected to an *in vitro* deubiquitination assay using ubiquitin-AMC. The fluorescence intensity of released AMC was measured. See Supplementary Fig. 2.

We also performed *in silico* homology modelling using the MAGI1 WW domain (PDB ID: 2YSE) [34] as a template. In this model, the overall structure of the autoinhibitory region of USP8 is similar to that of canonical WW domains, i.e. a three-stranded anti-parallel β-sheet [33] (Fig. 2**c**, left panel). Similar to WW domains, the β-sheet surface contains several aromatic or hydrophobic side chains (Phe^658^, Pro^661^, Phe^666^, Tyr^668^, His^670^ and His^677^) (Fig. 2**c**, right panel), indicating its role in protein–protein interactions. In addition, the N-terminal region, where Leu^651^ and Pro^652^ make contact with Trp^655^, forms a hook structure that is often found in WW domains. Molecular dynamics (MD) simulations showed the stability of the model structure; they also revealed that substitution of Trp^655^ to Ser (W655S) abolishes one of three β-strands and induces destabilisation of the domain structure (Fig. 2**d**). Thus, the first conserved tryptophan residue seems to underpin the folding, similar to the process in WW domains. Based on the common features of WW domains, we designated this region a WW-like domain.

We also examined the deubiquitinating activity of USP8 in which Trp^655^ is substituted to Ser (USP8^W655S^); we found that USP8^W655S^ shows higher activity than that of USP8^WT^ (Fig. 2**e**; Supplementary Figs. 2**a**, **b**). These results suggest that Trp^655^ in the WW-like domain is important for its inhibitory function as well as folding.

### The WW-like domain interacts with the catalytic USP domain

We hypothesised that the WW-like domain interacts with the USP domain. A pull-down assay indicated the interaction of aa 1–714 of USP8 (USP8^1–714^) with the USP domain, whereas deletion of the WW-like domain and substitution of Trp^655^ to Ser abolished the interaction (Figs. 3**a, b**; Supplementary Fig. 3**a**). These results indicate that the WW-like domain interacts with the USP domain *in vitro*.

**Fig. 3.**
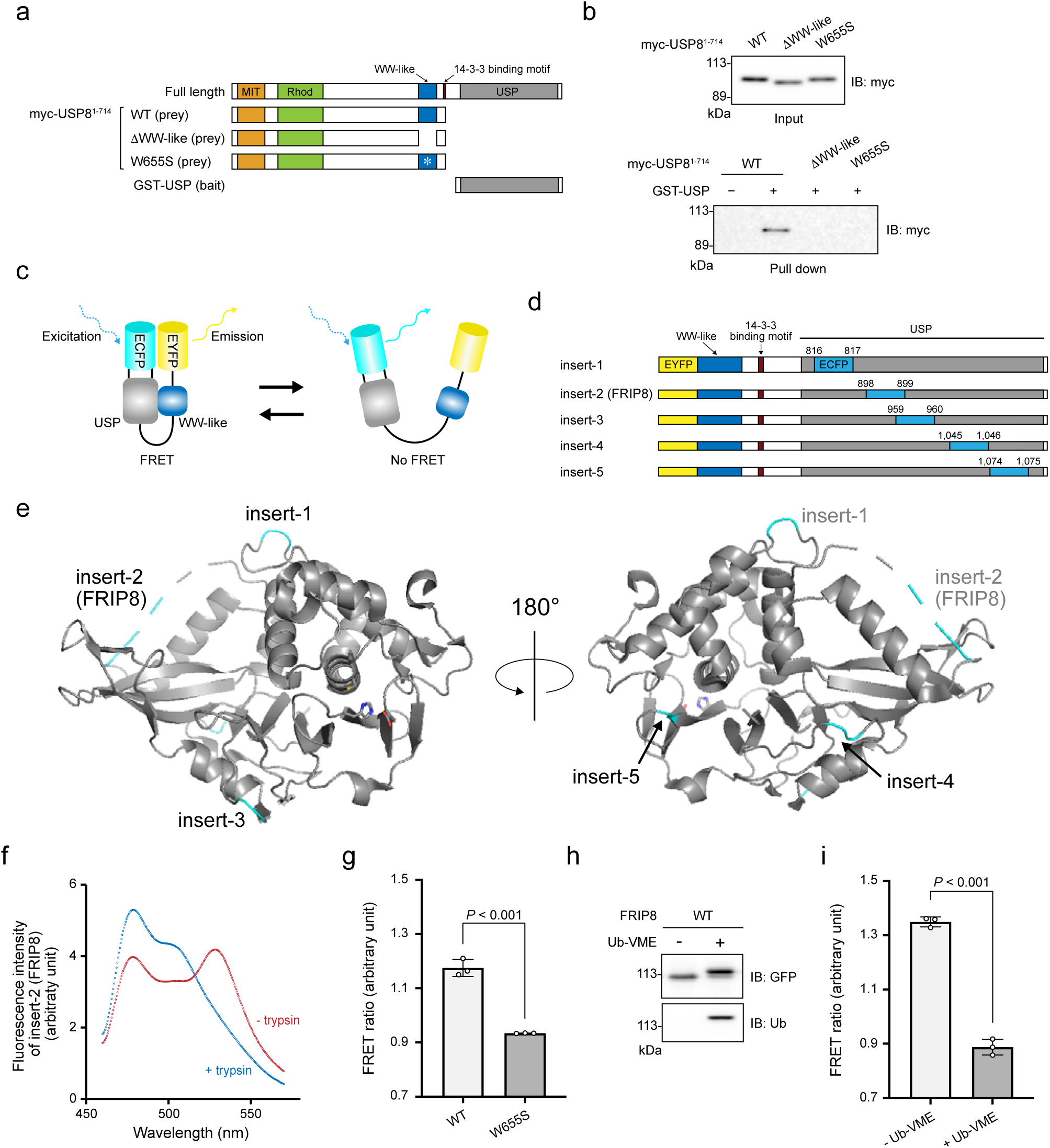
The interaction between the WW-like domain and the USP domain. **(a)** The schematic structures of the prey and bait proteins used in b. **(b)** Pull-down assay. Lysates of HEK293T cells expressing myc-tagged USP8^1–714^ (WT or mutants) were incubated with GST-tagged USP domain, subsequently precipitated using glutathione beads. The adsorbed fraction and input were analysed by immunoblotting. **(c)** Schematic representation of single-molecule FRET probes used to detect the interaction. **(d)** Schematic structures of the probes. **(e)** ECFP insertion points depicted on a crystal structure of the USP domain (PDB ID: 2GFO). Two representations are rotated by 180° around the Y-axis. **(f)** Emission spectra of FRIP8 at an excitation wavelength of 433 nm. Lysate of HEK293T cells expressing FRIP8 was subjected to fluorescence measurement with (blue) and without (red) trypsinisation. **(g)** FRET ratio of FRIP8^W655S^. The emission spectra of FRIP8^W655S^ were measured by the similar methods to those described in f. The ratio of the emission intensity of EYFP to that of ECFP (i.e. the FRET ratio) was calculated. The graph shows the means ± standard deviations of three independent experiments. Statistical significance was determined using Student’s *t*-test. **(h, i)** Ub-VME-labelling of FRIP8. Purified His-tagged FRIP8 was labelled with Ub-VME. It was subjected to immunoblotting (h) and a FRET assay (i). The graph shows the means ± standard deviations of three independent experiments. Statistical significance was determined using Student’s *t*-test. See Supplementary Fig. 3.

We also performed a single-molecule fluorescence resonance energy transfer (FRET) assay (Fig. 3**c**) by constructing five FRET probes as shown in Fig. 3**d**. These probes consist of USP8^645–1118^ (a region containing the WW-like domain and the USP domain), EYFP and ECFP. Specifically, EYFP was fused to the N-terminus of the WW-like domain, while ECFP was inserted into five different surface loops of the USP domain (Fig. 3**e**). Insertion of ECFP in these probes is thought not to disrupt the tertiary structure of the USP domain because many USP family proteins have long insertion sequences at these surface loops [19]. When the WW-like domain and USP domain interact in these probes, a fluorescent signal from EYFP can be detected in response to the excitation of ECFP.

We prepared lysates of cells expressing these probes and measured emission spectra with the excitation wavelength for ECFP. As negative controls, we trypsinised the lysates to separate EYFP and ECFP in the probes, and then we measured their emission spectra. All probes showed significant FRET signals (Supplementary Fig. 3**b**). The expression of each probe and their trypsinisation were evaluated by immunoblotting (Supplementary Fig. 3**c**). FRET signals of insert-2 and -3 probes, in which the ECFP insertion sites are close to the ubiquitin-binding pocket (Fig. 3**e**), were relatively higher. This implies that the pocket would be involved in the efficient interaction with the WW domain. We named the insert-2 probe the FRET-based intramolecular interaction probe of USP8 (FRIP8), which is the name used hereafter. The detailed FRET spectra of FRIP8 are shown in Fig. 3**f**.

W655S substitution in FRIP8 significantly decreased the FRET signal (Fig. 3**g**; Supplementary Fig. 3**d**), indicating that proper folding of the WW-like domain is important for the interaction with the USP domain. We also purified His-tagged FRIP8 (Supplementary Fig. 3**e**) and treated it with Ub-VME, which was conjugated to FRIP8 (Fig. 3**h**) and decreased the FRET signal (Fig. 3**i**), suggesting that the WW-like domain and ubiquitin molecule competitively interact with the ubiquitin-binding pocket.

A previous report indicated that USP8 dimerisation potentially occurs via the N-terminal MIT domain [20]. We found that USP8 dimerisation occurred by co-immunoprecipitation of FLAG-tagged and myc-tagged USP8, but deletion of neither the WW-like domain nor the USP domain abolished this dimerisation (Supplementary Fig. 3**f**). Thus, these domains are apparently not required for USP8 dimerisation. Given that our FRET probes lacked the MIT domain, we speculate that the WW-like domain and the USP domain interact in an intramolecular manner in our FRET probes.

### The WW-like domain plugs the catalytic cleft of the USP domain and narrows the entrance to the ubiquitin-binding pocket

In further tests, we analysed the interaction between the WW-like domain and USP domain *in silico*. First, we predicted the USP domain structure in ubiquitin-bound form by homology modelling using the ubiquitin-bound USP2 USP domain (PDB ID: 2HD5) [35] as a template. Unlike its apo-form (PDB ID: 2GFO) [20], the predicted structure showed an open conformation in which the USP domain grabs the ubiquitin molecule with the tip of the fingers and the catalytic cleft (Fig. 4**a**; Supplementary Fig. 4**a**). Lys^913^ and Leu^917^ in a *α*9 helix (aa 902–918) (Fig. 1**a**) also make physical contact with ubiquitin. The C-terminal tail of ubiquitin is embedded in the catalytic cleft, supported by BL1 and BL2. The catalytic triad is aligned in the model structure.

**Fig. 4.**
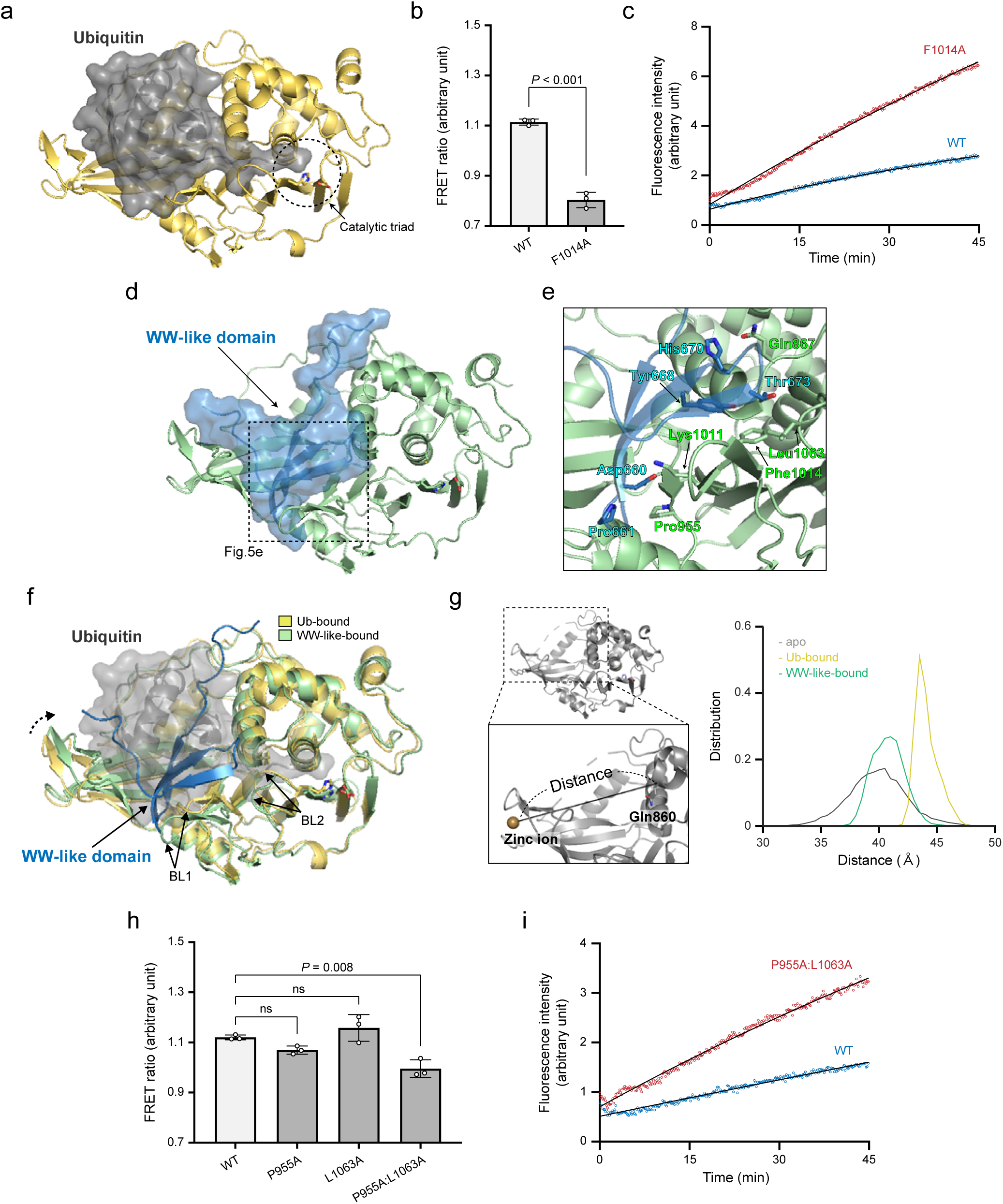
Structural modelling of the WW-like domain and USP domain complex. **(a)** Model structure of the USP8 USP domain in ubiquitin-bound form. The USP domain and the bound ubiquitin are shown in yellow and grey, respectively. **(b)** The FRET ratio of FRIP8^F1014A^, analysed by similar methods to those described in Fig. 3g. The graph shows the means ± standard deviations of three independent experiments. Statistical significance was determined using Student’s *t*-test. **(c)** Deubiquitination activities of USP8^F1014A^ toward ubiquitin-AMC, analysed by the similar methods to those described in Fig. 2e. **(d)** Complex structures of the WW-like domain (blue) and USP domain (green), as predicted by docking simulations using structural models of the WW-like domain in Fig. 2c and the USP domain in a. **(e)** Plugging of the catalytic cleft by the WW-like domain in the model. **(f)** Narrowing of the entrance to the ubiquitin-binding pocket caused by the interaction in the model. The USP domain in ubiquitin-bound form and that in WW-like domain-bound form are shown in yellow and green, respectively. **(g)** Distributions of the distance between zinc ions in the fingers subdomain and C*α* atoms of Gln^860^ during the MD simulations of the WW-like domain-bound form (green), ubiquitin-bound form (yellow) and apo-form (grey). **(h)** The FRET ratio of FRIP8^P955A^, FRIP8^L1063A^ or FRIP8^P955A:L1063A^, analysed by similar methods to those described Fig. 3g. The graph shows the means ± standard deviations of three independent experiments. Statistical significance against the WT was determined by one-way ANOVA and Tukey’s post hoc test. ns, not significant. **(i)** Deubiquitination activities of USP8^P955A:L1063A^ toward ubiquitin-AMC, analysed by the similar methods to those described in Fig. 2e. See Supplementary Fig. 4.

We also attempted to identify the putative binding site(s) by examining the effects of point mutations in FRIP8 on the FRET signal. Results showed that substitution of Phe^1014^ in the BL1 to Ala (F1014A) significantly decreased the FRET signal (Fig. 4**b**; Supplementary Fig. 4**b**). In addition, we examined the effects of F1014A mutation on the deubiquitinating activity of full-length USP8. Hereafter, we measured the activity using ubiquitin-AMC assay, which is more sensitive and quantitative than gel-based assay. The F1014A mutation enhanced the activity (Fig. 4**c**; Supplementary Fig. 4**c**). These findings suggest the involvement of Phe^1014^ in the binding process.

Using structural data from the WW-like domain (Fig. 2**c**) and USP domain (Fig. 4**a**), we performed *in silico* docking simulations. We selected the most probable structural model under the conditions that Phe^1014^ is one of the binding sites and that the inter-domain interaction is the most favourable possibility (Supplementary Fig. 4**d**; see Methods for more detail). We found that this stable complex model included a WW-like domain occupying part of the ubiquitin-binding pocket and plugging the catalytic cleft (Fig. 4**d**). An enlarged view of the contact sites is shown in Fig. 4**e**. Tyr^668^ on the β-sheet of the WW-like domain forms a hydrophobic contact with Phe^1014^ in the BL1. Thr^673^ in the third β-strand of the WW-like domain makes contact with Leu^1063^ in the BL2. Asp^660^ and His^670^ in the loops between β-strands in the WW-like domain form polar interactions with Lys^1011^ and Gln^867^ in the palm subdomain, respectively. BL1 and BL2 are likely relocated by these contacts from their positions in the ubiquitin-bound form (Fig. 4**f**).

Another important feature of this model is the slight bending of the fingers subdomain to the inside of the ubiquitin-binding pocket, which narrows the entrance to the pocket (Fig. 4**f**). Pro^661^ in the loop between the β-strands of the WW-like domain makes contact with Pro^955^ at the base of the fingers subdomain (Fig. 4**e**), which possibly contributes to the bending of the fingers. MD simulations confirmed that, although the fingers subdomain moves flexibly in the apo-form, it is fixed to this position in the WW-like domain-bound form (Fig. 4**g**).

We also conducted mutation analysis targeting two putative binding sites: Pro^955^ in the fingers subdomain and Leu^1063^ in the BL2. Combined substitution of Pro^955^ and Leu^1063^ to Ala (P955A:L1063A) decreased the FRIP8 FRET signal, although single amino acid substitutions with each residue separately had no effect (Fig. 4**h**; Supplementary Fig. 4**b**). Importantly, despite mutations in the ubiquitin-binding pocket, P955A:L1063A enhanced the deubiquitinating activity of full-length USP8 (Fig. 4**i**; Supplementary Fig. 4**e**). Taken together with effects of F1014A (Figs. 4**b**, **c**), these results clearly demonstrate a binding–inhibition relationship. Notably, Pro^955^ and Leu^1063^ as well as Phe^1014^ are highly conserved among vertebrate species (Supplementary Fig. 4**f**).

### 14-3-3 inhibits USP8 enzyme activity partly by enhancing the interaction between the WW-like domain and the USP domain

Studies have shown that 14-3-3 binds to USP8 in a phosphorylation-dependent manner [29]. We measured the 14-3-3 binding motif phosphorylation levels in total cell lysates and in absorbed fractions with GST-tagged 14-3-3; in the latter, all USP8 molecules should be phosphorylated. By comparing the results, we estimated that ∼60% of cellular USP8 is phosphorylated at the 14-3-3 binding motif (Supplementary Fig. 5**a**).

R18 is a peptide that inhibits 14-3-3 interaction with its ligands [36]. Consistent with previous reports, R18 treatment induced the dissociation of 14-3-3 from USP8 (Fig. 5**a**) and enhanced the deubiquitinating activity of USP8 immunoprecipitates (Fig. 5**b**; Supplementary Fig. 5**b**). FRIP8 harbours a 14-3-3 binding motif between the WW-like domain and the USP domain (Fig. 3**d**). Importantly, R18 treatment induced the dissociation of 14-3-3 from FRIP8 immunoprecipitates (Fig. 5**c**) and reduced the FRET signal (Fig. 5**d**). These results indicate that 14-3-3 enhances the interaction between the WW-like domain and USP domain.

**Fig. 5.**
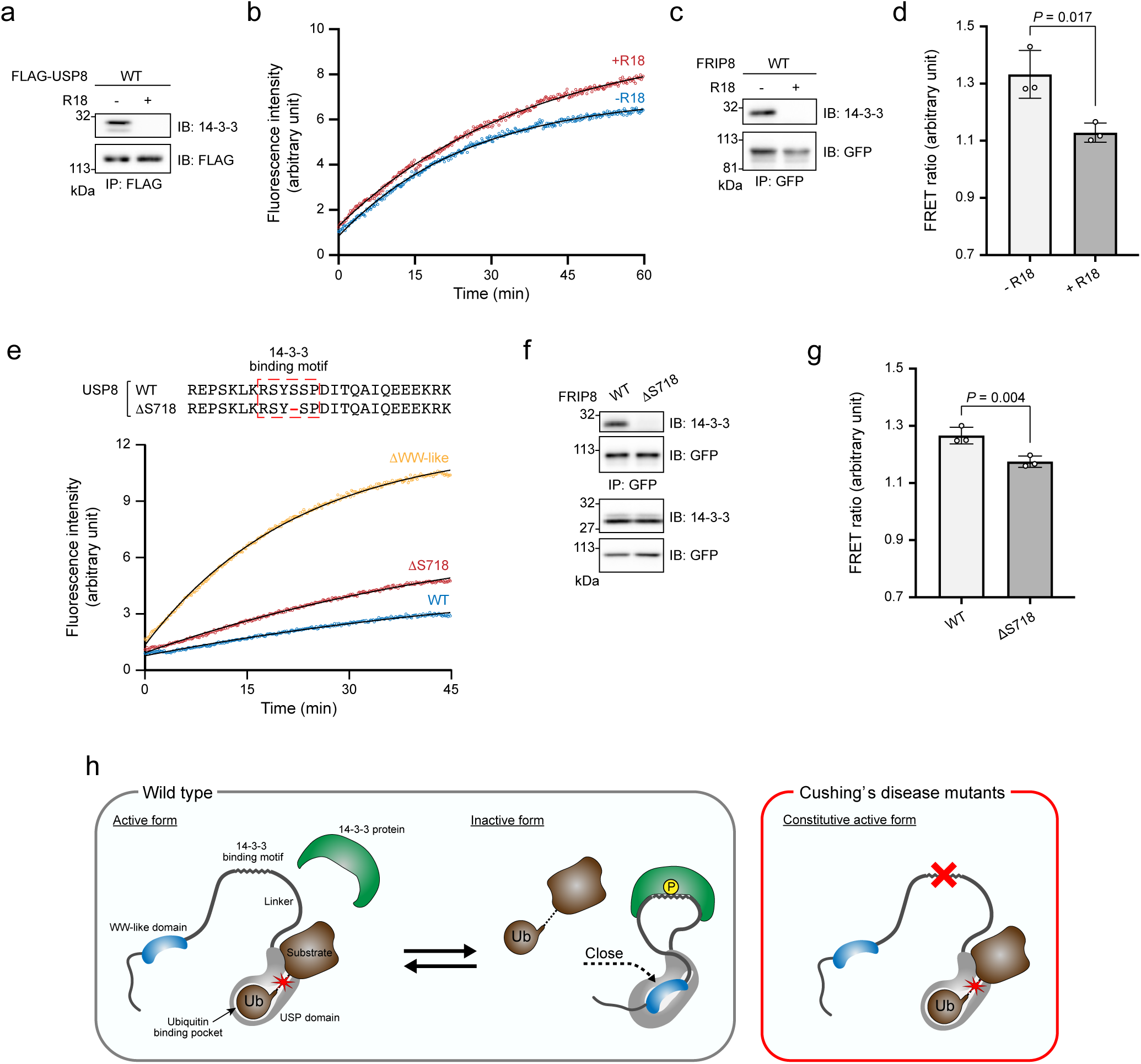
Effects of 14-3-3 on the interaction of the WW-like domain with the USP domain. **(a)** Effects of R18 on co-immunoprecipitation of USP8 with 14-3-3. HEK293 cells expressing FLAG-tagged USP8 were lysed. The lysate was incubated with 100-μM R18 before being subjected to immunoprecipitation and immunoblotting. **(b)** Effects of R18 on USP8 deubiquitinating activity. The lysates of HEK293 cells expressing FLAG-tagged USP8 were incubated with 100-μM R18 before they were subjected to an *in vitro* deubiquitination assay similar to that described in Fig. 2e. **(c, d)** Effects of R18 on FRIP8. HEK293 cells expressing FRIP8 and 14-3-3*ε* were lysed. The lysate was incubated with 100-μM R18 and then subjected to immunoprecipitation and subsequently immunoblotting (c). Samples were analysed using similar methods to those described in Fig. 3g (d). The graph shows the means ± standard deviations of three independent experiments. Statistical significance was determined using Student’s *t*-test. **(e)** Deubiquitination activities of USP8s with Cushing’s disease-associated mutations. The amino acid sequences of the mutants are shown in the top panel. The lysates of HEK293 cells expressing USP8 mutants were subjected to immunoprecipitation; the deubiquitinating activities in immunoprecipitates were then analysed using similar methods to those described in Fig. 2e. **(f, g)** Effects of Cushing’s disease-associated mutations on FRIP8. HEK293 cells expressing FRIP8 mutants were lysed and subjected to immunoprecipitation. Samples were then analysed by immunoblotting (f) and a FRET assay (g). The graph shows the means ± standard deviations of three independent experiments. Statistical significance against the WT was determined by one-way ANOVA and Tukey’s post hoc test. **(h)** The working hypothesis of the novel autoinhibitory mechanism of USP8 and its impairment in Cushing’s disease mutants.

Deletion of Ser^718^ (ΔS718) is a common USP8 mutation in Cushing’s disease that cause dissociation from 14-3-3 [30, 37]. A previous report indicated that Ser^718^ deletion enhances USP8 deubiquitinating activity [30]. We compared the effects of deleting Ser^718^ or the WW-like domain on USP8 deubiquitinating activity. Deletion of Ser^718^ enhanced this activity more weakly than did deletion of the WW-like domain (Fig. 5**e**; Supplementary Fig. 5**c**). Importantly, Ser^718^ deletion was found to induce the dissociation of 14-3-3 from FRIP8 immunoprecipitates (Fig. 5**f**) and to slightly reduce the FRET signal (Fig. 5**g**; Supplementary Fig. 5**d**). It indicates that Ser^718^ deletion significantly but incompletely inhibited the interaction between the WW-like domain and USP domain. Collectively, Ser^718^ deletion enhances USP8 deubiquitinating activity, partly by suppressing the interaction between the WW-like domain and USP domain.

## Discussion

In this study, we showed that the USP8 WW-like domain and ubiquitin competitively bind to the ubiquitin-binding pocket of USP8. In most WW domains, a few aromatic or hydrophobic side chains on the intrinsic β-sheet make contact with short hydrophobic sequences (e.g. Pro-Pro-X-Tyr) in ligand proteins [33, 38]. In contrast, our docking simulation highlights a distinct feature of the USP8 WW-like domain: several amino acids on the β-sheet and at the peripheral loops make contact with a large surface produced by the different subdomains of the USP domain. Similarly, ubiquitin binds to the USP domains through a large contact area [39]. Through this large contact, the WW-like domain appears to plug the catalytic cleft of the USP domain. In addition, the WW-like domain alters the position of the fingers subdomain, probably by contacting with Pro^955^ in this subdomain, and thereby narrows the entrance to the pocket. To our knowledge, USP8 is the only USP family protein with a WW-like domain, which suggests that regulation via the WW-like domain may be unique to USP8.

Several USPs are reported to be regulated by the intrinsic regions that physically interact with their catalytic domains [26, 28, 40–42]. Among these, USP25 had previously been the only USP for which the activity was inhibited on a well-defined structural basis. The present study provides another example of the autoinhibition of USPs by an intrinsic region. In USP25, the autoinhibitory region occupies a space between the *α*5 helix and BL2 [41, 42]. Interestingly, the autoinhibitory regions of USP8 and USP25 similarly plug an entrance to the catalytic cleft in which the ubiquitin C-terminal tail should be embedded. In a case where a USP was inhibited by a compound, a similar space in USP14 was occupied by the small molecule inhibitor IU1 [43]. Therefore, this space could potentially be utilised for effective inhibition of many USPs by intrinsic regulatory motifs and chemical inhibitors.

We found that the majority of cellular USP8 is bound to 14-3-3. Our analyses also indicated that 14-3-3 inhibits USP8 deubiquitinating activity by enhancing the interaction between the WW-like and USP domain. Since the 14-3-3-binding motif is located at an unstructured region between the WW-like domain and the USP domain, one possibility is that binding shortens the distance between these domains and thereby enhances their complex formation (Fig. 5**h**). However, further study is needed to elucidate the underlying mechanism.

The 14-3-3 binding motif is a hot spot at which somatic mutations frequently occur in Cushing’s disease [30]. This study revealed one possible part of the pathogenic mechanism underlying Cushing’s disease, i.e. the inability of USP8 to bind to 14-3-3 decreases the interaction between the WW-like and USP domains, which causes USP8 hyperactivation (Fig. 5**h**). To date, researchers of Cushing’s disease tumours have found various mutations around the 14-3-3 binding motif but none in the WW-like domain. Mutations in the WW-like domain may cause cell death rather than ACTH hypersecretion by enhancing USP8 deubiquitinating activity too strongly or by affecting the other functions of USP8. A previous study showed that mutations within the 14-3-3 binding motif cause the cleavage of USP8, thereby producing a 40-kDa C-terminal fragment (C40) with excess deubiquitination activity [30]. Surprisingly, we did not detect C40 in our study. Since different cell lines (Cos-7 cells vs. HEK293/HEK293T cells) were used in the two studies, C40 may be produced in a cell type-specific manner. It will therefore be important to determine whether significant levels of C40 are produced in the corticotrophs of Cushing’s disease patients. Nevertheless, our study demonstrated that Cushing’s disease-associated mutations release USP8 autoinhibition, which causes high enzymatic activity even without its cleavage.

In conclusion, we found that the USP8 WW-like domain functions as a novel autoinhibitory region by binding to the ubiquitin-binding pocket of the catalytic domain. We also found that 14-3-3 binds to USP8 and thereby enhances the autoinhibitory interaction. In cells, USP8 deubiquitinating activity seems to be tuned by the association and dissociation of 14-3-3. In Cushing’s disease, this regulation is apparently impaired by disease-associated mutations at the 14-3-3 binding motif. To build upon our findings, further studies will be required to determine the physiological and pathological significance of this new regulatory mechanism of USP8.

## Methods

### cDNA preparation and plasmid construction

Human USP8 cDNA [30], human 14-3-3ε cDNA [29], mouse STAM1 cDNA [44] and mouse STAM2 cDNA [45] were prepared elsewhere. Human Parkin cDNA and human ubiquitin cDNA were kindly donated by Dr. Koji Yamano and Dr. Toshiaki Suzuki, respectively (Tokyo Metropolitan Institute of Medical Science, Japan). For the construction of expression plasmids, these cDNAs were subcloned into the following vectors: pME [30] for the expression of C-terminal FLAG-tagged USP8; pCDH-CMV-MCS-EF1-Puro (System Biosciences) for the viral infection and expression of N-terminal FLAG-tagged USP8; pcDNA3 (Invitrogen) for N-terminal HA-tagged Ubiquitin; pFLAG-CMV2 (Sigma Aldrich) for N-terminal FLAG-tagged USP8, Parkin, STAM1 and STAM2; pmyc-CMV5 (gifted from Dr. Jun Nakae, Keio University, Tokyo, Japan) for N-terminal myc-tagged USP8 and 14-3-3ε; and pGEX-6P2 (GE Healthcare) for the bacterial expression of N-terminal GST-tagged USP domain (aa 756–1118).

For construction of single-molecule FRET probes, human USP8 (aa 645–684) was subcloned into pEYFP-N1 (Clonetech). ECFP with flexible spacers consisting of a 10-aa peptide (-Gly-Gly-Ser-Ala-Gly-Gly-Ser-Ala-Gly-Gly-) was amplified by PCR from pECFP-N1 (Clonetech) [46, 47]. For ECFP insertion, restriction sites for EcoRI and BamHI were inserted in the following points of the USP domain: gaps between aa 816/817 as insert-1, aa 898/899 as insert-2 (FRIP8), aa 959/960 as insert-3, aa 1045/1046 as insert-4 and aa 1074/1075 as insert-5 [19]. ECFP with the spacers was subcloned in a stepwise manner into these EcoRI and BamHI sites. For construction of His-tagged FRIP8, His-tag sequence was fused to the C-terminal of FRIP8. Site-specific mutagenesis was achieved using PCR with a Prime STAR Mutagenesis Basal Kit (Takara).

### Protein preparation

GST-tagged USP domain (aa 756–1118) was produced in *Escherichia coli* Rosetta. Protein expression was induced by the stimulation of cells with 0.5-mM IPTG, which was followed by overnight culture at 15°C. Cells were then lysed in ice-cold PBS supplemented with 1% Triton-X-100 and protease inhibitors (1-μg/ml aprotinin, 1-μg/ml leupeptin and 1-μg/ml pepstatin A). After centrifugation, the supernatants were mixed with glutathione sepharose 4B (GE Healthcare). After 1-h incubation at 4°C, beads were washed five times with PBS and then incubated in ice-cold elution buffer A [50-mM Tris-HCl, pH 8.0; 1-mM EDTA; 1-mM DTT; and 10-mM reduced glutathione (Wako)] at 4°C for 30 min. The purity and concentration of eluates were validated by SDS-PAGE and CBB staining using Quick CBB (Wako).

His-tagged FRIP8 was expressed in HEK293T cells. Cell culture and transfection are described below. Cells were lysed in ice-cold lysis buffer (50-mM Tris-HCl, pH 7.4; 150-mM NaCl; 50-mM NaF; and 1% Triton-X-100) supplemented with 10-mM imidazole and protease inhibitors. After centrifugation, the supernatants were mixed with Ni-NTA agarose beads (Qiagen). After 2-h incubation at 4°C, beads were washed five times with Tris-buffered saline (TBS) [20-mM Tris-HCl (pH 7.4) and 150-mM NaCl], and then incubated in ice-cold elution buffer B (20-mM Tris-HCl, pH 7.4; 150-mM NaCl; and 250-mM imidazole) at 4°C for 30 min. The purity was validated using immunoblotting.

For the *in vitro* deubiquitination assay and ubiquitin-VME labelling, FLAG-tagged USP8 (wild-type and mutants) were expressed in HEK293 cells (for ubiquitin-AMC assay) or HEK293T cells (for ubiquitin chain cleavage assay and for ubiquitin-VME labelling). Cell lysis and immunoprecipitation were carried out as described below. USP8 levels in each sample were compared by immunoblotting, with an equal number of USP8 molecules being used in the assay.

### Cell culture, plasmid transfection and lentivirus infection

HEK293, HEK293T and HeLa cells were grown in Dulbecco’s modified Eagle’s medium (DMEM; Nacalai Tesque) supplemented with 10% foetal bovine serum (FBS), 100-units/ml penicillin and 0.1-mg/ml streptomycin at 37°C and 5% CO_2_. To stimulate cells with EGF, they were cultured in the presence of 0.5% FBS for the last 24 h and then incubated with 100-ng/ml human EGF (PeproTech) at 37°C for 5 min.

Plasmid transfections were conducted using polyethyleneimine (Polyscience) according to the standard protocol. Cells were subjected to analyses 24–48 h after transfection.

For lentivirus production, HEK293T cells were transfected with pCDH-CMV-FLAG-USP8-EF1-Puro, psPAX2 (Addgene #12260) and pCMV-VSV-G (Addgene #8454). Two days after transfection, the medium containing the virus was collected and filtrated. HeLa cells were infected with the virus medium in the presence of 8-μg/ml polybrene (Nacalai Tesque). From 2 days after infection, cells were cultured in the presence of 0.8-µg/ml puromycin. Survived cells were used for experiments.

### Cell lysis, immunoprecipitation and immunoblotting

Cells were lysed with ice-cold lysis buffer (50-mM Tris-HCl, pH 7.4; 150-mM NaCl; 50-mM NaF and 1% Triton-X-100) supplemented with protein inhibitors. To detect ubiquitin signals by immunoblotting, 2-mM N-ethylmaleimide (Sigma Aldrich) was added. For the immunoprecipitation of FLAG-tagged USP8 for the deubiquitination assay, 1-mM DTT (Nacalai Tesque) was added. After centrifugation of the lysates, the supernatants were collected. To examine effects of R18 peptide (Sigma Aldrich) on the interaction of 14-3-3 and USP8 or FRIP8, the lysates were incubated with 100-μM R18 peptide at 4°C for 30 min.

Immunoprecipitation was performed using standard procedures. Anti-FLAG M2 antibody-conjugated agarose beads (Sigma Aldrich), anti-EGFR antibody (#MI-12-1, MBL), anti-GFP antibody (#M-048-3, MBL) and Protein A-Sepharose (GE Healthcare) were used for immunoprecipitation. After immunoprecipitation, beads were washed five times with lysis buffer. For the elution of FLAG-tagged proteins, beads were incubated in TBS with 200-ng/μl FLAG peptide (Sigma Aldrich) at 4°C for 30 min.

Samples were incubated in SDS-PAGE sample buffer (62.5-mM Tris-HCl, pH 6.8; 2% SDS; 5% 2-mercapto ethanol; 10% glycerol; and 0.1-mg/ml bromo phenol blue) at 98°C for 5 min. Otherwise, for the detection of ubiquitin signals, samples were incubated in NuPAGE LDS sample buffer with reducing agent (Thermo Scientific) at 37°C for 20 min. Samples were separated by SDS-PAGE or NuPAGE (Thermo Scientific).

Immunoblotting was performed using standard procedures. The primary antibodies used for immunoblotting were as follows: anti-FLAG antibody (clone 1E6, Wako), anti-Ubiquitin (#D058-3, MBL; #3936, Cell Signalling), anti-EGFR antibody (#MI-12-1, MBL), anti-USP8 antibody [48], anti-HRS antibody [49], anti-STAM1 antibody [50], anti-α-tubulin antibody (clone 10G10, Wako), anti-HA antibody (clone 3F10, Sigma Aldrich; #SC-57592, Santa Cruz), anti-Myc antibody (#SC-40, Santa Cruz; clone 9E10, hybridoma supernatant), anti-GFP antibody (#A-11122, Thermo Scientific) and anti-14-3-3 antibody (#SC-1657, Santa Cruz). The secondary antibodies used were peroxidase-conjugated anti-mouse IgG, anti-rat IgG and anti-rabbit IgG antibodies (GE Healthcare). Blots were detected using ECL Prime Western Blotting Detection Reagents (GE Healthcare) and ImageQuant LAS 4000 Mini (GE Healthcare).

### *In vitro* deubiquitination assay

For ubiquitin chain cleavage assay, USP8 (wild-type or the mutants) was incubated with 25 ng/μl of Lys63-linked ubiquitin tetramers (Boston Biochem) in TBS with 10-mM dithiothreitol (DTT) at 37°C for the indicated times in the presence or absence of R18 peptide. Samples were subjected to SDS-PAGE or NuPAGE and then to gel staining with a Silver Stain MS Kit (Wako).

For the ubiquitin-AMC assay, USP8 was mixed with 1-μM ubiquitin-AMC (Boston Biochem or Life Sensors) and 10-mM DTT in TBS in 96-well medium-binding, flat-bottom, black plates (Thermo Scientific). To examine the effects of R18 peptide, USP8 was incubated with 100-μM R18 peptide at 4°C for 30 min before the assay. Plates were set on Varioskan LUX (Thermo Scientific), and incubated at 37°C. During the incubation, the fluorescence intensity of AMC released from ubiquitin was measured at 10-s intervals using 345 nm and 445 nm as the excitation and emission wavelengths, respectively. The fluorescence intensity of the sample without USP8 was measured as the background and then subtracted from each value. Non-linear regression curves were plotted using the equation for one-phase association by Prism 9 (GraphPad Software).

### Ubiquitin-VME labelling

USP8 immunoprecipitates or His-tagged FRIP8 precipitates were incubated with 1-μM ubiquitin-VME (Boston Biochem) in TBS supplemented with 1-mM DTT at 37°C for 5 min or 30 min, respectively. Samples were then subjected to immunoblotting and a FRET assay.

### Immunofluorescence

HeLa cells on coverslips were fixed with PBS containing 4 % paraformaldehyde, permeabilised with PBS containing 0.2 % Triton X-100, and blocked with PBS containing 5 % FBS. Cells were stained with mouse anti-FLAG antibody (Sigma Aldrich) and Alexa Fluor 488-conjugated anti-mouse IgG secondary antibody (Invitrogen), according to the standard procedures. Coverslips were mounted on slides with Fluoroshield mounting medium (ImmunoBioScience). Fluorescence images were captured with a laser-scanning confocal microscope (LSM 780, Carl Zeiss).

### Amino acid sequence alignment

Protein BLAST (https://blast.ncbi.nlm.nih.gov/) was used to identify protein sequences similar to USP8 aa 645–684. Sequence similarity was analysed using ClustalW (https://clustalw.ddbj.nig.ac.jp/); the colours in the alignment followed the ClustalX scheme. Amino acid sequences of human MAGI1 (accession code: Q96QZ7), human SAV1 (Q9H4B6), human WWOX (Q9NZC7), and vertebrate USP8s were based on annotated information in the UniProt database (https://www.uniprot.org/).

### *In silico* structural modelling

*In silico* structural modelling of the WW-like domain, the USP domain, and their complex was performed. Briefly, (1) homology modelling of the WW-like domain was first conducted using the MAGI1 WW domain (PDB ID: 2YSE), (2) the structural model of the USP domain in the ubiquitin-bound form was then constructed using the ubiquitin-bound USP2 USP domain (PDB ID: 2HD5) as the template for homology modelling, and (3) the protein–protein docking simulations of the two domains were executed using ClusPro (https://cluspro.org/). The MD simulations of the derived modelled structures were also performed to examine their stabilities in the physiological condition; the details are described as follows.

By use of the sequency similarity with MAGI1 WW domain (see Fig. 2b) together with the structural data by NMR experiment (PDB ID: 2YSE), the structure of WW-like domain in USP8 (aa 645–684) was modelled by MODELLER [51]. To perform the MD simulation, a rectangular simulation box was constructed with a margin of 12 Å to the boundary of the simulation box, resulting in the dimension, 58 Å × 51 Å × 58 Å. The solution system contained 4139 TIP3P water molecules [52] together with one sodium ion to neutralize the simulation system, resulting in 13045 atoms in total. AMBER ff14SB [53] was applied for the potential energy of the all-atom protein. For the zinc ion in the fingers subdomain, the Zinc AMBER force field (ZAFF) [54, 55] was used. The MD simulations were performed by AMBER 16 [56] under constant temperature and pressure (NPT) conditions at *T* = 300 K and *P* = 1 atm using Berendsen’s thermostat and barostat [57] at a relaxation time of 1 ps, and using the particle mesh Ewald method [58] for the electrostatic interactions. The simulation length was 1 μs, together with a 2-fs time step using constraining bonds involving hydrogen atoms via the SHAKE algorithm [59]. The simulation of the W655S mutant was also carried out as described above after replacing W655 of the wild type with Ser via MODELLER [51].

The structural model of the USP domain (aa 756–1110) was also constructed to survey the potential interaction with the WW-like domain. Since the open form with bound ubiquitin was not solved for USP8, the ubiquitin-bound USP2 USP domain (PDB ID: 2HD5) was used as the template of the homology modelling. The modelled structure of the USP8 USP domain with bound 76-residue ubiquitin was then solvated in the simulation box of the dimension, 103 Å × 84 Å × 91 Å (including 65415 atoms in total), and simulated for 1 μs as described above. The stability of the modelled ubiquitin-USP domain complex was seen in the small root mean square distribution on C*α* atom resolution (C*α*-RMSD) of the overall structure being < 2.6 Å from the initial simulation model. The structure of the USP domain in the last simulation trajectory time step was then used for the subsequent docking simulation with the WW-like domain. The MD simulation of apo USP domain in the closed form was also carried out by use of the crystal structure (PDB ID: 2GFO) as the initial model.

Structural basis of stable complex formation between the USP domain and the WW-like domain was then pursued by use of their docking simulation. The protein-protein docking simulation was performed using protein-protein interaction server ClusPro together with the standard parameter sets [60], generating putative interaction complexes. Among them, we chose six model candidates from the criteria that F1014 of the USP domain exists in the interface with the WW-like domain, and carried out 100-ns MD simulation for each model candidate as described above. Using the associated MD trajectories, the number of atom contacts (non-hydrogen atom pair with *r* < 4 Å), *N*_c_, and binding free energy evaluated using MM-GB/SA module of AMBER 16 [56], ΔG_bind_, were calculated between the USP domain and the WW-like domain. The structure with the largest *N*_c_ and the lowest ΔG_bind_ was then selected as the most probable complex model (see Supplementary Fig. 4d). The additional 1-μs MD simulation of the resulting USP domain/WW-like domain complex showed that the overall C*α*-RMSD is less than 3.5 Å from the initial simulation model, indicating substantial stability of the derived complex structure in physiological condition.

### Pull-down assay

HEK293T cells expressing indicated prey proteins were lysed with lysis buffer supplemented with protein inhibitors. The lysates were incubated with 5-μM GST or GST-tagged USP domain at 4°C for 2 h, and then incubated in the presence of glutathione sepharose 4B at 4°C for 90 min. Beads were then washed five times with lysis buffer and samples were subjected to immunoblotting.

### FRET assay

Samples were added to 96-well medium-binding, flat-bottom, black plates. Fluorescence intensities were measured using Varioskan LUX or F2700 (Hitachi Hightech) at 5-nm or 0.5-nm wavelength intervals, respectively, with 433 nm as the excitation wavelength and 460–570 nm as the emission wavelength range. Negative control samples were incubated with 10 ng/μl of trypsin (Thermo Scientific) at 4°C for 60 min, and then fluorescence intensities were measured. To examine the effects of R18 peptide, samples were incubated with 100-μM R18 peptide at 4°C for 30 min before the assay. In blank wells, assay buffer or lysates derived from cells that did not express FRET probes were also prepared. These fluorescence intensities were measured as background signals and subtracted from measured values. The FRET ratio was calculated using the peak intensity at the EYFP emission wavelength divided by the peak intensity at the ECFP emission wavelength.

### Statistical analysis

The results are shown as the mean ± standard deviation. Data were analysed using Student’s t-tests or one-way factorial ANOVA followed by Tukey–Kramer multiple comparison post hoc tests. P values <0.05 were considered to show statistical significance.

### Uncropped scan data

Uncropped scan data of blots and staining are shown in Supplementary Figs. 6, 7.

## Acknowledgements

We thank Dr. Sayaka Yasuda (Tokyo Metropolitan Institute of Medical Science) for helpful discussions and Dr. Koji Yamano (Tokyo Metropolitan Institute of Medical Science) for the kind donation of Parkin plasmids. We also acknowledge the help of the Biomaterials Analysis Division (Tokyo Institute of Technology) with DNA sequencing analysis. This work was supported by JSPS KAKENHI Grant Number 17K08625, 19H05289 and 21H00276 to T.F.; JSPS KAKENHI Grant Number 15H04293 to M. Komada; Platform Project for Supporting Drug Discovery and Life Science Research [Basis for Supporting Innovative Drug Discovery and Life Science Research (BINDS)] from AMED under Grant Number JP21am0101109 to M.I.; and the Nagase Science and Technology Foundation.

## Abbreviations

ACTH: adrenocorticotropic hormone
BL: blocking loop
C40: C-terminal 40 kDa fragment of USP8
CBB: coomassie brilliant blue
CHMP: charged multivesicular body protein
COPII: coat protein complex II
DMEM: Dulbecco’s modified Eagle medium
DTT: DL-dithiothreitol
DUSP: domain in ubiquitin-specific protease
ECFP: enhanced cyan fluorescence protein
EDTA: ethylenediaminetetraacetic acid
EGFR: epidermal growth factor receptor
EPG5: ectopic P granules protein 5 homolog
Eps15: epidermal growth factor receptor substrate 15
EYFP: enhanced yellow fluorescence protein
FBS: foetal bovine serum
FRET: fluorescence resonance energy transfer
FRIP8: FRET-based intramolecular interaction probe of USP8
GST: glutathione S-transferase
HRS: hepatocyte growth factor-regulated tyrosine kinase substrate
MAGI1: membrane-associated guanylate kinase, WW and PDZ domain-containing protein 1
MD: molecular dynamics
MIT: microtubule interacting and transport domain
Ni-NTA: nickel nitrilotriacetic acid
PBS: phosphate buffered saline
PDB: protein data bank
Rhod: rhodanese-like domain
SAV1: protein salvador homolog 1
SBM: SH3 binding motif
SNARE: soluble NSF attachment protein receptor
STAM: signal transducing adaptor molecule
TBS: Tris-buffered saline
Ub-AMC: ubiquitin-conjugated 7-amino-4-methylcoumarin
Ub-VME: ubiquitin-conjugated vinyl methyl ester
Ub: ubiquitin
UBL: ubiquitin-like domain
USP: ubiquitin-specific protease
VAMP8: vesicle-associated membrane protein 8
WWOX: WW domain-containing oxidoreductase

## References

1. Mizuno, E. et al. Regulation of epidermal growth factor receptor down-regulation by UBPY-mediated deubiquitination at endosomes. Mol Biol Cell 16, 5163–5174 (2005).

2. Niendorf, S. et al. Essential role of ubiquitin-specific protease 8 for receptor tyrosine kinase stability and endocytic trafficking in vivo. Mol Cell Biol 27, 5029–5039 (2007).

3. Le Clorennec, C. et al. The anti-HER3 (ErbB3) therapeutic antibody 9F7-F11 induces HER3 ubiquitination and degradation in tumors through JNK1/2- dependent ITCH/AIP4 activation. Oncotarget 7, 37013–37029 (2016).

4. Oh, Y.M. et al. USP8 modulates ubiquitination of LRIG1 for Met degradation. Sci Rep 4, 4980 (2014).

5. Li, S. et al. Hedgehog-regulated ubiquitination controls smoothened trafficking and cell surface expression in Drosophila. PLoS Biol 10, e1001239 (2012).

6. Ma, G. et al. Regulation of Smoothened Trafficking and Hedgehog Signaling by the SUMO Pathway. Dev Cell 39, 438–451 (2016).

7. Xia, R., Jia, H., Fan, J., Liu, Y. & Jia, J. USP8 promotes smoothened signaling by preventing its ubiquitination and changing its subcellular localization. PLoS Biol 10, e1001238 (2012).

8. Mukai, A. et al. Balanced ubiquitylation and deubiquitylation of Frizzled regulate cellular responsiveness to Wg/Wnt. EMBO J 29, 2114–2125 (2010).

9. Mizuno, E., Kobayashi, K., Yamamoto, A., Kitamura, N. & Komada, M. A deubiquitinating enzyme UBPY regulates the level of protein ubiquitination on endosomes. Traffic 7, 1017–1031 (2006).

10. Row, P.E., Prior, I.A., McCullough, J., Clague, M.J. & Urbe, S. The ubiquitin isopeptidase UBPY regulates endosomal ubiquitin dynamics and is essential for receptor down-regulation. J Biol Chem 281, 12618–12624 (2006).

11. Crespo-Yanez, X. et al. CHMP1B is a target of USP8/UBPY regulated by ubiquitin during endocytosis. PLoS Genet 14, e1007456 (2018).

12. Adoro, S. et al. Post-translational control of T cell development by the ESCRT protein CHMP5. Nat Immunol 18, 780–790 (2017).

13. Gu, H. et al. USP8 maintains embryonic stem cell stemness via deubiquitination of EPG5. Nat Commun 10, 1465 (2019).

14. Peng, H. et al. The ubiquitin-specific protease USP8 directly deubiquitinates SQSTM1/p62 to suppress its autophagic activity. Autophagy 16, 698–708 (2020).

15. Durcan, T.M. et al. USP8 regulates mitophagy by removing K6-linked ubiquitin conjugates from parkin. EMBO J 33, 2473–2491 (2014).

16. Kawaguchi, K. et al. Ubiquitin-specific protease 8 deubiquitinates Sec31A and decreases large COPII carriers and collagen IV secretion. Biochem Biophys Res Commun 499, 635–641 (2018).

17. Mukai, A. et al. Dynamic regulation of ubiquitylation and deubiquitylation at the central spindle during cytokinesis. J Cell Sci 121, 1325–1333 (2008).

18. Clague, M.J., Urbe, S. & Komander, D. Breaking the chains: deubiquitylating enzyme specificity begets function. Nat Rev Mol Cell Biol 20, 338–352 (2019).

19. Ye, Y., Scheel, H., Hofmann, K. & Komander, D. Dissection of USP catalytic domains reveals five common insertion points. Mol Biosyst 5, 1797–1808 (2009).

20. Avvakumov, G.V. et al. Amino-terminal dimerization, NRDP1-rhodanese interaction, and inhibited catalytic domain conformation of the ubiquitin-specific protease 8 (USP8). J Biol Chem 281, 38061–38070 (2006).

21. Kohler, A., Zimmerman, E., Schneider, M., Hurt, E. & Zheng, N. Structural basis for assembly and activation of the heterotetrameric SAGA histone H2B deubiquitinase module. Cell 141, 606–617 (2010).

22. Li, H. et al. Allosteric Activation of Ubiquitin-Specific Proteases by beta-Propeller Proteins UAF1 and WDR20. Mol Cell 63, 249–260 (2016).

23. Yin, J. et al. Structural Insights into WD-Repeat 48 Activation of Ubiquitin-Specific Protease 46. Structure 23, 2043–2054 (2015).

24. Schlicher, L. et al. SPATA2 promotes CYLD activity and regulates TNF-induced NF-kappaB signaling and cell death. EMBO Rep 17, 1485–1497 (2016).

25. Wagner, S.A., Satpathy, S., Beli, P. & Choudhary, C. SPATA2 links CYLD to the TNF-alpha receptor signaling complex and modulates the receptor signaling outcomes. EMBO J 35, 1868–1884 (2016).

26. Faesen, A.C. et al. Mechanism of USP7/HAUSP activation by its C-terminal ubiquitin-like domain and allosteric regulation by GMP-synthetase. Mol Cell 44, 147–159 (2011).

27. Hu, M. et al. Structure and mechanisms of the proteasome-associated deubiquitinating enzyme USP14. EMBO J 24, 3747–3756 (2005).

28. Xue, W. et al. Domain interactions reveal auto-inhibition of the deubiquitinating enzyme USP19 and its activation by HSP90 in the modulation of huntingtin aggregation. Biochem J 477, 4295–4312 (2020).

29. Mizuno, E., Kitamura, N. & Komada, M. 14-3-3-dependent inhibition of the deubiquitinating activity of UBPY and its cancellation in the M phase. Exp Cell Res 313, 3624–3634 (2007).

30. Reincke, M. et al. Mutations in the deubiquitinase gene USP8 cause Cushing’s disease. Nat Genet 47, 31–38 (2015).

31. Ma, Z.Y. et al. Recurrent gain-of-function USP8 mutations in Cushing’s disease. Cell Res 25, 306–317 (2015).

32. Borodovsky, A. et al. Chemistry-based functional proteomics reveals novel members of the deubiquitinating enzyme family. Chem Biol 9, 1149–1159 (2002).

33. Salah, Z., Alian, A. & Aqeilan, R.I. WW domain-containing proteins: retrospectives and the future. Front Biosci (Landmark Ed) 17, 331–348 (2012).

34. Ohnishi, S. et al. Structural basis for controlling the dimerization and stability of the WW domains of an atypical subfamily. Protein Sci 17, 1531–1541 (2008).

35. Renatus, M. et al. Structural basis of ubiquitin recognition by the deubiquitinating protease USP2. Structure 14, 1293–1302 (2006).

36. Petosa, C. et al. 14-3-3zeta binds a phosphorylated Raf peptide and an unphosphorylated peptide via its conserved amphipathic groove. J Biol Chem 273, 16305–16310 (1998).

37. Perez-Rivas, L.G. et al. The Gene of the Ubiquitin-Specific Protease 8 Is Frequently Mutated in Adenomas Causing Cushing’s Disease. J Clin Endocrinol Metab 100, E997–1004 (2015).

38. Lin, Z., Xie, R., Guan, K. & Zhang, M. A WW Tandem-Mediated Dimerization Mode of SAV1 Essential for Hippo Signaling. Cell Rep 32, 108118 (2020).

39. Ernst, A. et al. A strategy for modulation of enzymes in the ubiquitin system. Science 339, 590–595 (2013).

40. Clerici, M., Luna-Vargas, M.P., Faesen, A.C. & Sixma, T.K. The DUSP-Ubl domain of USP4 enhances its catalytic efficiency by promoting ubiquitin exchange. Nat Commun 5, 5399 (2014).

41. Sauer, F. et al. Differential Oligomerization of the Deubiquitinases USP25 and USP28 Regulates Their Activities. Mol Cell 74, 421–435 e410 (2019).

42. Gersch, M. et al. Distinct USP25 and USP28 Oligomerization States Regulate Deubiquitinating Activity. Mol Cell 74, 436–451 e437 (2019).

43. Wang, Y. et al. Small molecule inhibitors reveal allosteric regulation of USP14 via steric blockade. Cell Res 28, 1186–1194 (2018).

44. Mizuno, E., Kawahata, K., Kato, M., Kitamura, N. & Komada, M. STAM proteins bind ubiquitinated proteins on the early endosome via the VHS domain and ubiquitin-interacting motif. Mol Biol Cell 14, 3675–3689 (2003).

45. Takata, H., Kato, M., Denda, K. & Kitamura, N. A hrs binding protein having a Src homology 3 domain is involved in intracellular degradation of growth factors and their receptors. Genes Cells 5, 57–69 (2000).

46. Levskaya, A., Weiner, O.D., Lim, W.A. & Voigt, C.A. Spatiotemporal control of cell signalling using a light-switchable protein interaction. Nature 461, 997–1001 (2009).

47. Komatsu, N. et al. Development of an optimized backbone of FRET biosensors for kinases and GTPases. Mol Biol Cell 22, 4647–4656 (2011).

48. Kato, M., Miyazawa, K. & Kitamura, N. A deubiquitinating enzyme UBPY interacts with the Src homology 3 domain of Hrs-binding protein via a novel binding motif PX(V/I)(D/N)RXXKP. J Biol Chem 275, 37481–37487 (2000).

49. Komada, M. & Kitamura, N. Growth factor-induced tyrosine phosphorylation of Hrs, a novel 115-kilodalton protein with a structurally conserved putative zinc finger domain. Mol Cell Biol 15, 6213–6221 (1995).

50. Mizuno, E., Kawahata, K., Okamoto, A., Kitamura, N. & Komada, M. Association with Hrs is required for the early endosomal localization, stability, and function of STAM. J Biochem 135, 385–396 (2004).

51. Sali, A. & Blundell, T.L. Comparative Protein Modeling by Satisfaction of Spatial Restraints. Journal of Molecular Biology 234, 779–815 (1993).

52. Jorgensen, W.D. Comparison of simple potential functions for simulating liquid water. J. Chem. Phys. 79, 926–935 (1983).

53. Maier, J.A. et al. ff14SB: Improving the Accuracy of Protein Side Chain and Backbone Parameters from ff99SB. J Chem Theory Comput 11, 3696–3713 (2015).

54. Peters, M.B. et al. Structural Survey of Zinc-Containing Proteins and Development of the Zinc AMBER Force Field (ZAFF). J Chem Theory Comput 6, 2935–2947 (2010).

55. Li, P.F., Roberts, B.P., Chakravorty, D.K. & Merz, K.M. Rational Design of Particle Mesh Ewald Compatible Lennard-Jones Parameters for +2 Metal Cations in Explicit Solvent. J Chem Theory Comput 9, 2733–2748 (2013).

56. Case, D.A. et al. The Amber biomolecular simulation programs. Journal of Computational Chemistry 26, 1668–1688 (2005).

57. Juffer, A.H., Botta, E.F.F., Vankeulen, B.A.M., Vanderploeg, A. & Berendsen, H.J.C. The electric potential of a macromolecule in a solvent: a fundamental approach. J. Comp. Phys. 97, 144–171 (1991).

58. Darden, T., York, D. & Pedersen, L. Particle Mesh Ewald - an N.Log(N) Method for Ewald Sums in Large Systems. Journal of Chemical Physics 98, 10089–10092 (1993).

59. Ryckaert, J.P., Ciccotti, G. & Berendsen, H.J.C. Numerical integration of the cartesian equations of motion of a system with constraints: molecular dynamics of n-alkanes. J. Comput. Phys. 23, 327–341 (1977).

60. Kozakov, D. et al. The ClusPro web server for protein-protein docking. Nat Protoc 12, 255–278 (2017).

