## Supplementary Figs. 1-7 for "The molecular basis of ubiquitin-specific protease 8 autoinhibition by the WW-like domain"

### Supplementary Fig. 1

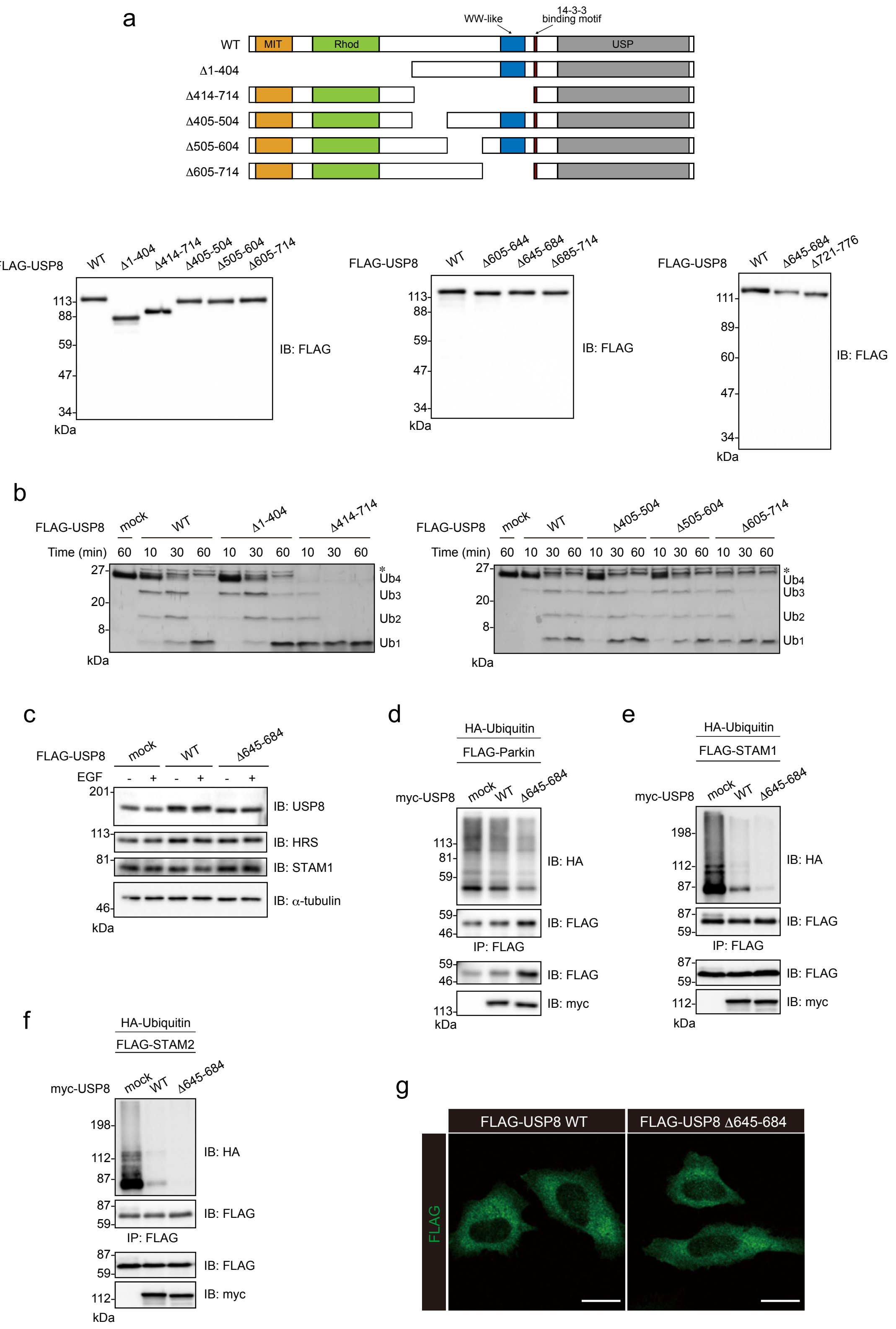

**Supplementary Fig. 1 (related to Fig. 1)**

(a) Top, schematic structures of USP8 mutants used in Supplementary Fig. 1b. Bottom, immunoblotting of anti-FLAG immunoprecipitates used in Fig. 1c and Supplementary Fig. 1b.

(b) Deubiquitination activities of indicated USP8 mutants, analyzed by the similar experiment to Fig. 1c. *, co-purified protein(s) with USP8.

(c) Immunoblotting analysis of cell lysates used in Fig. 1e, confirming similar expression levels of exogenous and endogenous USP8.

(d) (e) (f) Parkin (d), STAM1 (e) and STAM2 (f) ubiquitination levels in cells overexpressing $USP8^{\Delta 645\text{-}684}$. HEK293T cells overexpressing myc-tagged USP8 (WT or Δ645-684), FLAG-tagged substrate proteins (Parkin, STAM1 or STAM2), and HA-tagged ubiquitin were treated with EGF. The lysates were subjected to immunoprecipitation and immunoblotting.

(g) Immunofluorescence of $USP8^{\Delta 645\text{-}684}$. HeLa cells expressing FLAG-tagged USP8 (WT or Δ645-684) were fixed and stained with anti-FLAG antibody for confocal microscopy. Scale bars, 20 μm.

#### Supplementary Fig. 2

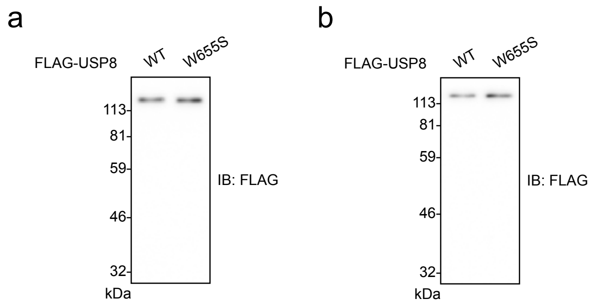


**Supplementary Fig. 2 (related to Fig. 2)**

(a) Immunoblotting of anti-FLAG immunoprecipitates used in Fig. 2e, top.

(b) Immunoblotting of anti-FLAG immunoprecipitates used in Fig. 2e, bottom.

Supplementary Fig. 3

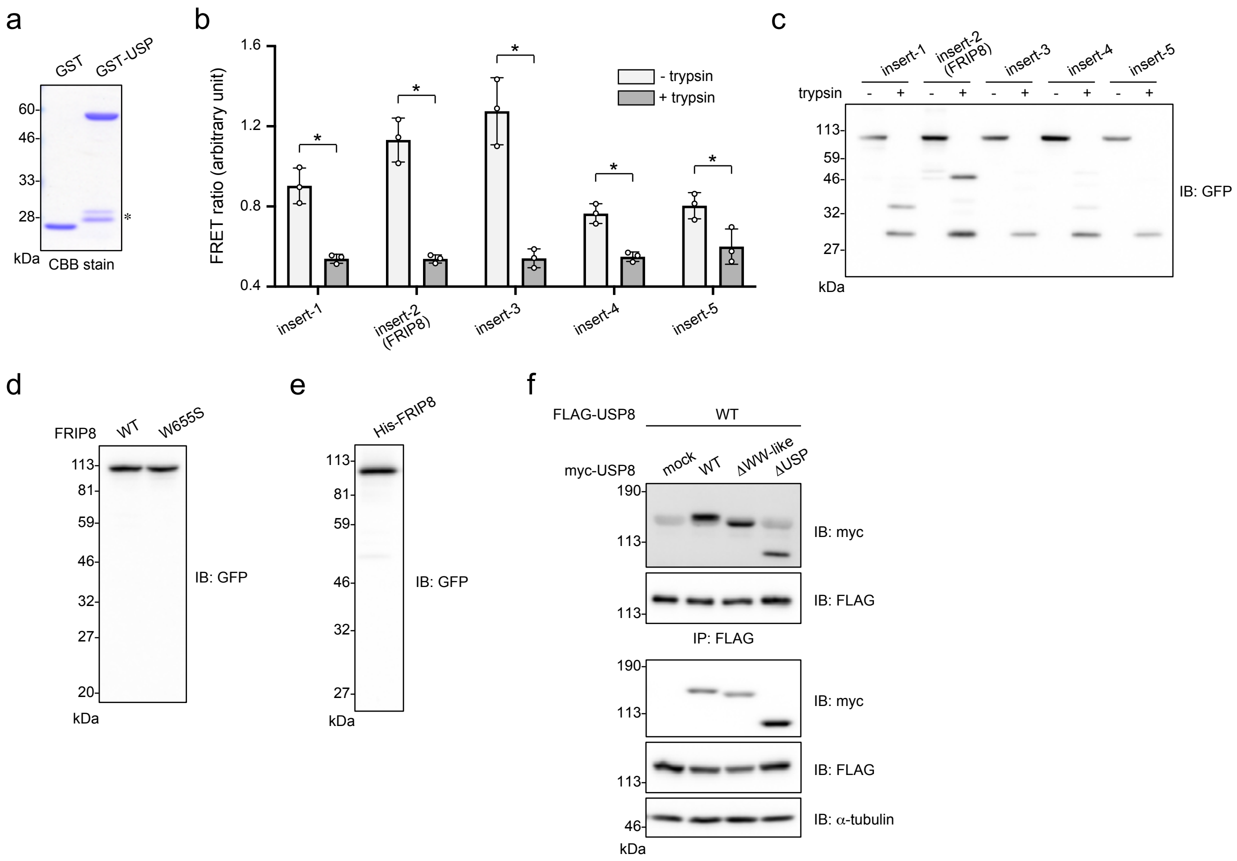


**Supplementary Fig. 3 (related to Fig. 3)**

(a) CBB stain of GST-tagged protein used in Fig. 3b. *, degradation products of GST-USP.

(b) FRET ratio of probes indicated in Fig. 3d. Lysates of HEK293T cells expressing each probe were subjected to fluorescence measurement before (white) and after (black) trypsinization. The ratio of the emission intensity of EYFP to that of ECFP (FRET ratio) was calculated. The graph shows the means ± SD of three independent experiments. Statistical significance was determined by Student's *t*-test. *, $P < 0.05$.

(c) Immunoblotting of cell lysates used in b, confirming proper expression and trypsin-dependent cleavage of the probes.

(d) Immunoblotting of FRIP8 (WT and W655S) in Fig. 3g. HEK293T cells expressing FRIP8 (WT and W655S) were lysed, and the lysates were subjected to immunoblotting.

(e) Immunoblotting of His-tagged FRIP8 used in Fig. 3f. HEK293T cells expressing His-tagged FRIP8 were lysed. FRIP8 was purified using Ni-NTA agarose, and subjected to immunoblotting.

(f) Roles of the WW-like domain and the USP domain in USP8 dimerization. HEK293T cells expressing indicated proteins were lysed, and the lysates were subjected to immunoprecipitation and immunoblotting.

### Supplementary Fig. 4

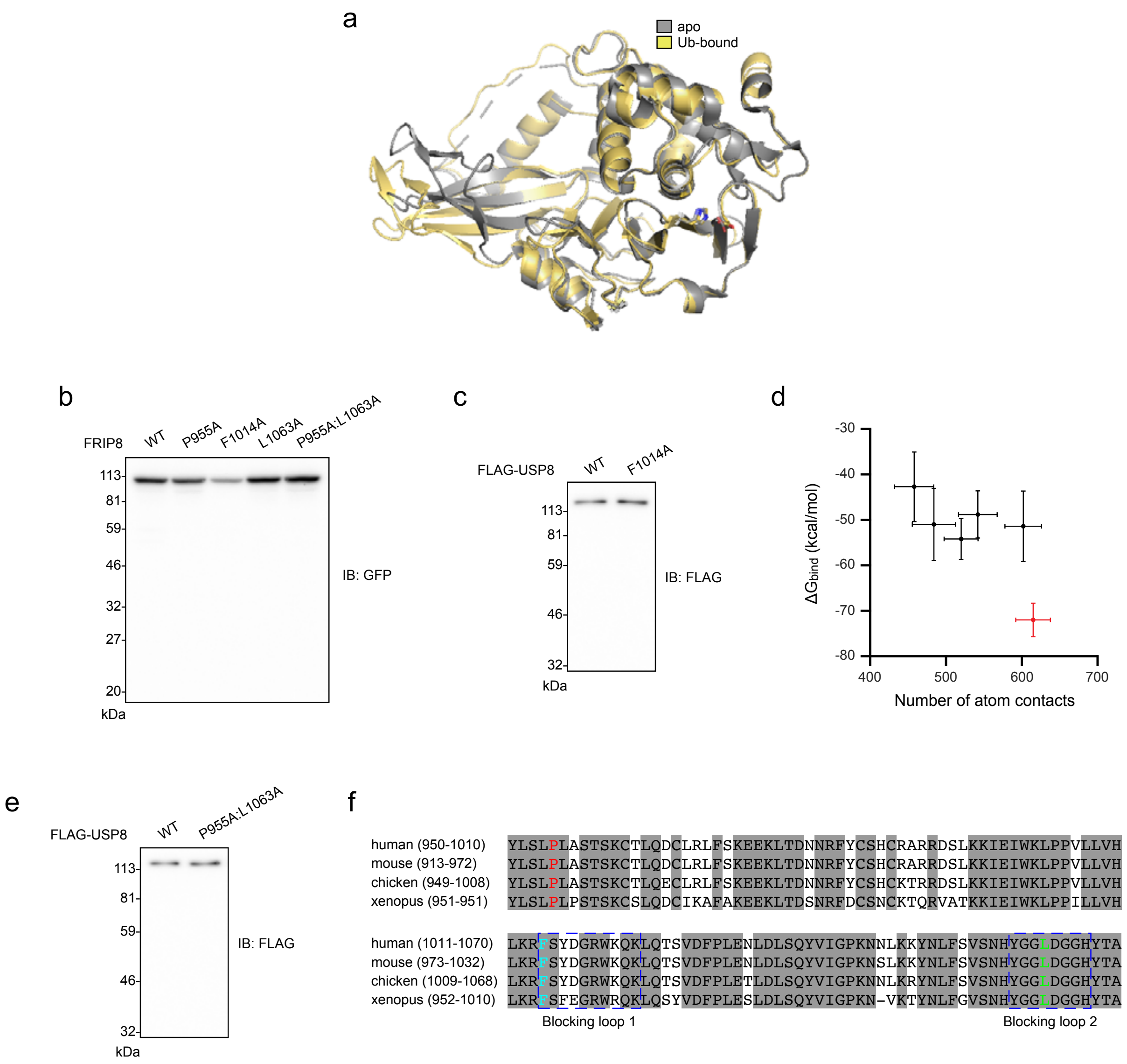

**Supplementary Fig. 4 (related to Fig. 4)**

(a) Superposition of the two USP domain structures, the ubiquitin-bound form of the present model (yellow: Fig. 4a) and the apo form (grey: PDB ID: 2GFO).

(b) Immunoblotting of FRIP8 mutants in Fig. 4b and 4h. HEK293T cells expressing FRIP8 mutants were lysed, and the lysates were subjected to immunoblotting.

(c) Immunoblotting of USP8 (WT or F1014A) used in Fig. 4c. HEK293 cells expressing FLAG-tagged USP8 (WT or F1014A) were lysed, and anti-FLAG immunoprecipitates were analyzed by immunoblotting.

(d) Number of atom contacts, $N_c$, and binding free energy, $\Delta G_{bind}$, between the USP domain and the WW-like domain are plotted for six model candidates. The selected model structure with the largest $N_c$ and the lowest $\Delta G_{bind}$ is shown by red. See Methods for definition.

(e) Immunoblotting of USP8 (WT or P955A:L1063A) used in Fig. 4i, by the similar analysis to c.

(f) Conservation of amino acid residues corresponding to human USP8 $Pro^{955}$ (red), $Phe^{1014}$ (blue) and $Leu^{1063}$ (green) in vertebrate USP8s.

Supplementary Fig. 5

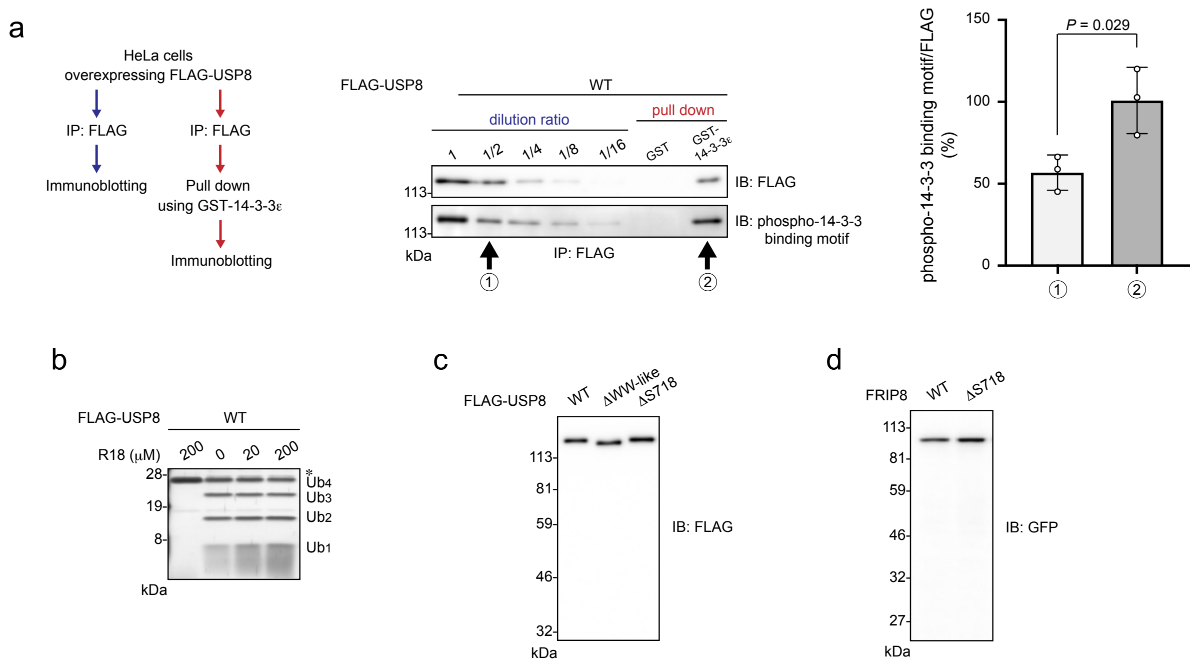


**Supplementary Fig. 5 (related to Fig. 5)**

(a) Percentage of USP8 14-3-3 binding motif phosphorylated in cells. Left panel, experimental flow. HeLa cells expressing FLAG-tagged USP8 were lysed, and anti-FLAG immunoprecipitates were diluted at various ratios. A portion of immunoprecipitates were incubated with GST-tagged 14-3-3 protein immobilized on glutathione beads. Samples were analyzed by immunoblotting with anti-FLAG antibody and anti-phospho-14-3-3 binding motif antibody. Arrows indicate lanes where the same amount of USP8 was loaded. Phosphorylation levels of 14-3-3 binding motif were measured by densitometric analyses. Right graph shows the means ± SD of three independent experiments. Statistical significance was determined by Student's *t*-test.

(b) Effects of R18 on USP8 activity. The lysates of HEK293T cells expressing FLAG-tagged USP8 were incubated with the indicated concentration of R18 before they were subjected to an *in vitro* deubiquitination assay similar to that described in Fig. 1c. *, co-purified protein(s) with USP8.

(c) Immunoblotting of USP8 mutants used in Fig. 5e, by the similar analysis to Supplementary Fig. 2e.

(d) Immunoblotting of FRIP8 mutants in Fig. 5g. HEK293 cells expressing FRIP8 mutants were lysed, and the lysates were subjected to immunoblotting.

### Supplementary Fig. 6

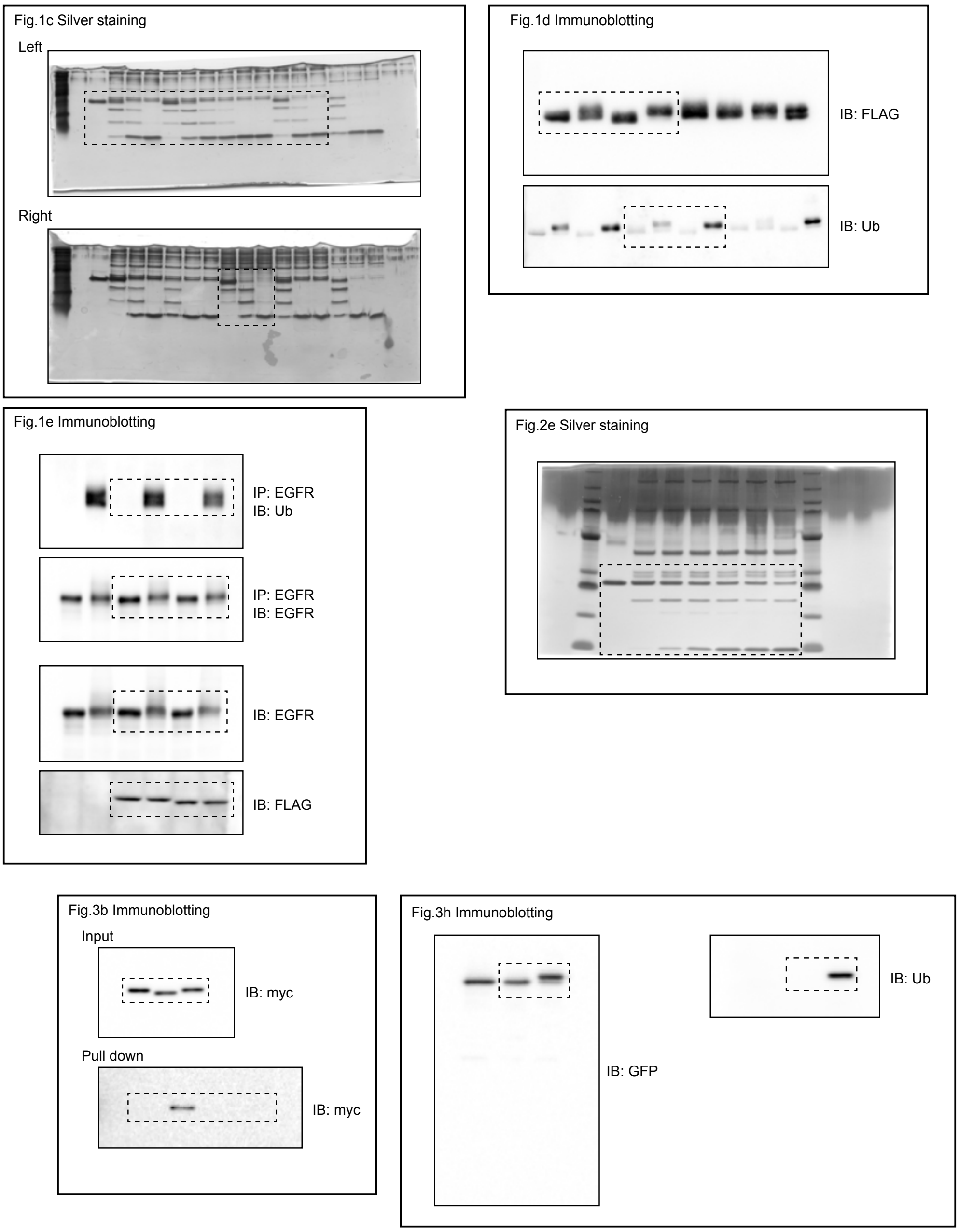


**Supplementary Fig. 6 Uncropped scans shown in Figs. 1c-e; Fig. 2e; Figs. 3b, h**

Dotted lined delimit cropped area used in the indicated panels.

#### Supplementary Fig. 7

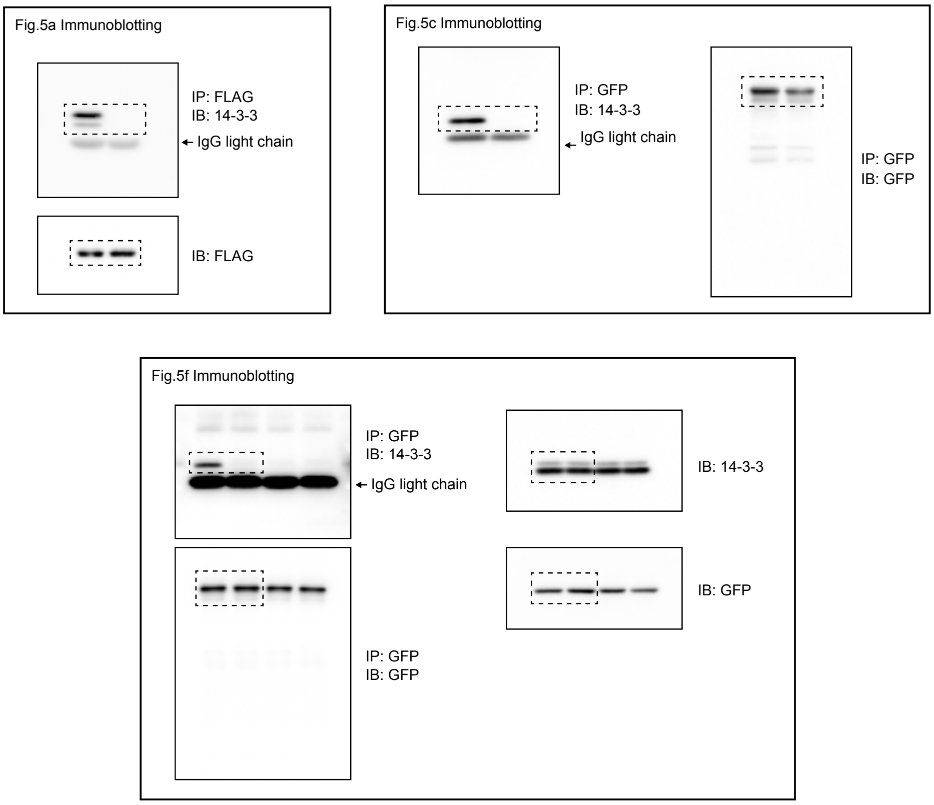


**Supplementary Fig. 7 Uncropped scans shown in Figs. 5a, c, f**

Dotted lined delimit cropped area used in the indicated panels.
